# Toxicology assessment of manganese oxide nanomaterials with enhanced electrochemical properties using human *in vitro* models representing different exposure routes

**DOI:** 10.1101/2022.04.30.490128

**Authors:** Natalia Fernández-Pampin, Juan José González Plaza, Alejandra García, Elisa Peña, Carlos Rumbo, Rocío Barros, Sonia Martel, Santiago Aparicio, Juan Antonio Tamayo-Ramos

## Abstract

In the present study, a comparative human toxicity assessment between newly developed Mn_3_O_4_ nanoparticles with enhanced electrochemical properties (GNA35) and their precursor material (Mn_3_O_4_) was performed, employing different *in vitro* cellular models representing alveolar, oral and dermal exposure routes, namely the human alveolar carcinoma epithelial cell line (A549), the human colorectal adenocarcinoma cell line (HT29), and the reconstructed 3D human epidermal model EpiDerm™ (RhE). The obtained results showed that Mn_3_O_4_ and GNA35 harbour similar morphological characteristics, while differences were observed in relation to their thermal stability and electrochemical properties. In regard to their toxicological properties, both nanomaterials induced oxidative stress in the A549 and HT29 cell lines that have demonstrated that manganese oxide nanoparticles reduce the cell viability in A549 cells while they do not produce any negative effect on that of HT29 cells. On the other hand, it was observed that Mn_3_O_4_ and GNA35 induce oxidative stress in both cell lines. Finally, it was noticed that none of the nanoparticles caused a reduction of the viability on the skin tissue so that they could not be classified as irritants. Our findings demonstrate that the evaluation of the toxicity of nanoparticles using different models is a critical aspect to increase the knowledge on their potential impact on human health.

## 1. Introduction

Manganese is one of the most abundant metals in the planet (Nádaská et al., 2010), mainly found in nature as MnO_2_ or Mn_3_O_4_). (Post, 1999), and a critical raw material in a number of applications, including the global steel industry, the production of aluminium alloys, and the development of Mn batteries (Clarke and Upson, 2017). In the last two decades, manganese oxide nanomaterials have become relevant compounds as well in a variety of areas, such as the development of energy storage devices (lithium-ion batteries, capacitors), biomedical applications, catalysis, etc., due to their inherent divergence in redox properties, morphology, crystalline structure, and surface nanoarchitectures (Dawadi et al., 2020; Ghosh et al., 2020). Specifically, Mn_3_O_4_ nanoparticles are of particular interest for the development of supercapacitators, due to their single black manganese structure at room temperature, good stability and controllable microstructure (Zhu et al., 2020). Within this particular field, efforts are being done to improve the characteristics of Mn_3_O_4_ based electrodes, e.g. to enhance the specific surface area and the conductivity of the nanomaterial (Peng et al., 2010; Zhu et al., 2020).

Besides its industrial relevance, manganese is an essential micronutrient needed for plant growth and for maintaining animals’ health and well-being. Due to its central role for the organism, Mn levels have to be kept finely regulated in order to keep homeostatic concentrations of this ion (Martinez-Finley et al., 2013). While lack of this element causes detrimental effects over the organism, excess impairs the proper functioning of the cell. One of the basal problems is the reactivity of the ion itself. Manganese can act over endogenous H_2_O_2_ and catalyse the release of ROS, such as ·OH through Fenton type of reactions (Wang et al., 2019). The accumulation of ROS can lead to apoptotic events (Circu and Aw, 2010) further compromising the stability of tissues and subsequently affect the functionality of entire organs (Singh et al., 2019). Choi *et al.* have shown that MnOx nanoparticles can induce ROS in human lung adenocarcinoma cells (Choi et al., 2010). Moreover, Frick *et al.* have reported than Mn_3_O_4_ interacts with gluthatione (GSH) and it can induce apoptosis in rat type II epithelial cells (Frick et al., 2011).

The increase of manganese oxide applications, particularly in nanomaterial form, may consequently lead to an increase in the human exposure and risk, due to their potential toxicity. These risks are especially evident at the manufacturer’s side where workers are subjected to occupational exposure (miners, smelters, welders and workers in dry-cell battery factories). Inhalation, ingestion (mainly via mucocilliary clearance and swallowing of respiratory secretions) and dermal contact, are the main routes of nanomaterials exposure (Alhadlaq et al., 2019). From those, the principal way of exposure occurs usually via inhalation, being size-dependent the distribution of the inhaled Mn nanoparticles in the respiratory system. A significant fraction of inhaled Mn nanoparticles can bypass the liver to enter the blood stream, from where they can bypass the blood-brain barrier via the olfactory tract (Roels et al., 1997; Lucchini et al., 2012; Zoni et al., 2012). Brain possesses a high permeability to manganese, and the accumulation of this metal in the brain can induce an irreversible neurological syndrome like Parkinson’s disease, which is common in workers who have been exposed to Mn (Takeda, 2003).

For the above-mentioned reasons, there is a need to evaluate the possible associated hazards of novel Mn_3_O_4_ nanomaterials. Results obtained from nanotoxicology assays can be as well an important instrument for the industry, in the development of safer formulations. In the current study, a novel engineered Mn_3_O_4_ nanomaterial, namely GNA35, and its precursor material (Mn_3_O_4_), have been characterized at physicochemical and toxicological level, employing three cellular models resembling the main routes of nanomaterials exposure: inhalation (A549 cell line), ingestion (HT29 cell line) and skin contact (reconstructed human epidermal model EpiDerm™).

## 2. Material and methods

### 2.1 Cell lines

The human alveolar carcinoma epithelial cell line A549 (ATCC, CCL-185) and human colorectal adenocarcinoma cell line HT29 (Sigma Aldrich) were utilized for toxicological evaluation. A549 cells were cultured in Dulbecco’s Modified Eagle’s Medium (DMEM) supplemented with 10% fetal bovine serum (FBS) and 1% penicillin-streptomycin solution. HT29 cells were cultured in McCoy’s 5A medium supplemented with 10% fetal bovine serum, 1% penicillin-streptomycin solution and 1% glutamine. Both cell lines were maintained at 37 °C in a humidified atmosphere containing 5% CO_2_.

The reconstructed human epidermal model EpiDerm™ was purchased from MatTek Life Sciences (MA, USA). The tissues were kept following the manufactures instructions (MatTek In Vitro Life Science Laboratories, 2020).

### 2.2 Nanomaterials

Trimanganese tetraoxide (Mn_3_O_4_ and manganese oxide (GNA35) were kindly provided by GNANOMAT (Madrid, Spain). Original stocks had a concentration of 4 mg mL^-1^ and were subdivided in smaller aliquots for homogeneity in the results. Prior to use, unique aliquots were sonicated in a water bath for 20 minutes. Sonicated aliquots were used once in the course of the experiments, and then discarded.

### 2.4 Synthesis and characterization of the manganese oxide nanoparticles

GNA35 (Mn_3_O_4_ nanoparticles) was synthetized through a novel nanotechnological process, based on patented procedures (Patent number ES2678419A1), using commercial Mn_3_O_4_ as precursor (Strem Chemicals, MA, USA). The method consists of the precipitation in a basic medium of metallic oxides obtained by the previous dissolution of precursor (60 mg) in 40 mL of dissolvent, which is a mixture 1:6 of malonic acid and water. During this process, the dissolution was sonicated in a bath for 1 h until it was dissolved, and the temperature and pH variations were controlled.

DLS and the *ζ*-potential determination was done using a Zetasizer Nano ZS90 (Malvern Instruments). The nanomaterials suspensions (20 μg/mL) were sonicated for 10 minutes prior to the analysis. In case of DLS determination, samples were diluted 10 times (1:10) prior to analysis. *ζ*-potential was measured using the M3-PALS method. The TEM analysis was performed using a JEOL 1011 high-resolution (HR) TEM. Samples were deposited on standard copper grids. Characterization by thermogravimetric analysis (TGA) was done using a thermo-gravimetric analyzer (METTLER TOLEDO TGA-2 Start system) from 40 °C to 1050 °C with an air flow-rate of 100 mL min^-1^ and a heating-rate of 10 °C min^-1^. The chemical composition and crystalline phase of the nanoparticles were determined by X-ray diffraction analysis, using X-Ray Polycrystalline Diffraction technique. The X-Ray power diffraction was recorded on a Panalytical XPERT Pro system equipped with Cu K radiation (λ = 0.15418 nm). The porosity of both nanomaterials was determined by analyzing their nitrogen adsorption-desorption isotherms, employing a Micromeritics ASAP 2420 V2.09 equipment. An Arbin (BT-G-5-10A) potentiostat was used for the electrochemical evaluation of the materials (Mn_3_O_4_ and GNA35). Cyclic voltam-metry analyses were recorder at 5 mV s-1 in a potential range of −0.8 to 0.3V and −0.6 to 0.4V for Mn_3_O_4_ and GNA35, respectively. Galvanostatic charge-discharge analyses were recorded at 1 A g^-1^ to estimate the specific capacitance (F g^-1^) for each material. Fabrication of the electrodes was done mixing Mn_3_O_4_ or GNA35 (80%) with carbon black (10%) and poly(vinylidene fluoride) (10%) using a agate mortar, followed by the addition of ethanol to make a homogeneous paste. Nickel foam was used as current collector. The electrodes were dried at 90 °C overnight. The electrochemical measurements were carried out in a 3-electrode cell using previously manufactured electrodes as working electrodes, and platinum wire and Ag/AgCl as counter and reference electrodes, respectively. A solution of 1M KOH was employed as electrolyte.

### 2.7 Toxicology assays

#### 2.7.1 MTT assay

Plates for MTT assays were seeded with 5 × 10^3^ cells per well (A549 cell line), or 1 × 10^4^ cells per well (HT29) and left for incubation at 37 °C for 24 h at 5% CO_2_ environment. After the first stage of incubation, media was removed. Assayed concentrations were added on the corresponding wells, where untreated cells received cultivation media, and negative control wells received sterile H_2_O. All solutions were prepared in the corresponding growth media but lowering the FBS to 1% (v/v) and without modification of the antibiotics. Incubation lasted for 24 h at continuous 37 °C, 5% CO_2_ environment. After 24 h exposure, media with nanomaterials were retrieved and wells washed with sterile PBS. A hundred μL of growth media supplemented with 0.5 mg/mL MTT were added per well, and after 3 h incubation it was removed. The resulting formazan crystals were solubilized with 100 μL DMSO, incubated for 15 min. at 37 °C in darkness, under continuously shaking. Absorbance was measured at 590 nm with a microplate reader (BioTek Synergy HT). Three replicates per dose were included in each experiment and at least two independent experiments were performed.

#### 2.7.2 Oxidative stress assay

We followed a reactive oxygen species (ROS) assay method adapted from Domi *et al.* (Domi et al., 2020), using a quantitative method for measuring ROS upon exposure to nanomaterials. In brief, A549 and HT29 cells were seeded in 96 micro-well plates, at a density of 3 × 10^4^ cells per well. After 24 h incubation, growth media was removed and washed once with 1x Hank’s Balanced Salt Solution (HBSS) pH 7.2 (NaOH adjusted), then incubated with 50 μM 2,7-dichlorofluorescein diacetate (DCFH-DA) for 30 min at 37 °C in darkness, with the exception of three wells incubated only with HBSS without dye. Upon incubation, we removed the buffer with dye and washed cells once with HBSS. Previously prepared nanomaterials dissolved in HBSS were added to each well at different concentrations ranging from 10 to 80 mg L^-1^, while 20 μM H_2_O_2_ served as a positive control. Exposure volume was always 200 μL. Two group of wells, one containing the cells previously not exposed to dye, and an additional second group, were exposed to HBSS corrected with H2O (the same amount contained in the nanoparticle suspension). We measured fluorescence in a microplate reader (BioTek Synergy HT, excitation wavelength, 485/20; emission wavelength 520/20) after 0, 30, and 60 min of exposition.

#### 2.7.3 *In Vitro* EpiDerm™ Skin Irritation Test

These assays were performed according to the *in vitro* EpiDerm™ Skin Irritation Test (EPI-200-SIT) protocol (MatTek In Vitro Life Science Laboratories, 2020). The reconstructed human epidermal model EpiDerm™ (EPI-200-SIT, MatTek, Bratislava) consists of normal human-derived epidermal keratinocytes cultured to form a multilayered highly differentiated model of the human epidermis. EpiDerm™ includes organized basal cells, spinous and granular layers, and a multilayered stratum corneum containing intercellular lamellar lipid layers arranged in patterns analogous to those found *in vivo*.

Upon receipt, the EpiDerm™ tissues were inspected for damage according to manufacture instructions and they were transferred to 6-well plates prefilled with 0.9 mL of assay medium (EPI-100-NMM provided with EpiDerm™ tissues) and incubated for 1 h. At the end of this pre-incubation, tissues were transferred to fresh media and incubated overnight. After the overnight incubation to release transport-stress, the tissues were topically exposed to the test materials for 60 minutes. Three tissues were used per test material (TC) and controls. The negative control consisted in tissues exposed to phosphate buffered saline (PBS) while in the positive control, tissues were exposed to a 5% solution of SDS in water. After exposure, tissues were rinsed with DPBS, blotted on an absorbent pad, and dried with a sterile cotton-tipped swab to remove the test substances from the surface of the tissue. After 24 h of incubation, the medium was collected for analysis of cytokines and then, the tissues were incubated for another 18 h in fresh medium. Finally, they were transferred to fresh medium during 42 h.

Cell viability after material exposure was determined using the MTT viability assay. A 1 mg mL^-1^ solution of MTT (included in the kit (EPI-200-SIT, MatTek, Bratislava)) in assay medium was prepared. Following exposure and rinsing, the tissues were placed in a 24-well plate containing 0.3 mL per well of the MTT medium and incubated for 3 h. After MTT incubation, the blue formazan salt was extracted with 2 mL per tissue of isopropanol. The plates were set on orbital shaker for 2 h at room temperature. The optical density at 570 nm was measured on a spectrophotometer (BioTek Synergy HT). Isopropanol alone was used as a blank. The percent viability of each tissue was calculated relative to negative control using the following equation:

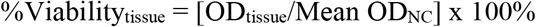

#### 2.7.4 Cytokine release assay

After treatment with Mn_3_O_4_, and GNA35 nanoparticles, culture media from the RhE tissues was recollected and stored at −20 °C. The concentration of IL-1α was evaluated by ELISA test (Diaclone), following the manufacturer’s instructions. Results are expressed as pg mL ^-1^ of cytokines released in the media.

#### 2.7.5 Statistical analysis

Statistical analysis data are represented as mean ± SD. Differences between the different treatments were established using a one-way ANOVA followed by multiple comparisons test (Tukey test). Statistical test was carried out using GraphPad Prim Software, Inc (version 8.0.2). Statistical significance was considered at P ≤ 0.05.

## 3. Results

### 3.1 Nanomaterials physicochemical characterization

GNA35 nanoparticles were synthesized aiming to obtain a nanomaterial with improved characteristics, such as surface area and conductivity (Peng et al., 2010; Zhu et al., 2020), for the development of supercapacitators. This was done using commercial Mn_3_O_4_ as precursor (Mn_3_O_4_ following an innovative procedure as described in Materials and Methods. Prior to their toxicological assessment, the physicochemical properties of the precursor material (Mn_3_O_4_ and the synthesized nanomaterial (GNA35) were determined, using several methodologies to obtain insights into their morphology, size, surface charge, crystallinity, surface area and electrochemical performance. Firstly, a TEM analysis was performed to observe their morphological characteristics and size. Figure 1 shows representative images of Mn_3_O_4_ (Figure 1a) and GNA35 (Figure 1b). As it can be observed, both the precursor and the synthesized materials showed to have round shape and a similar diameter, of around 100 nm or smaller.

**Figure 1.**
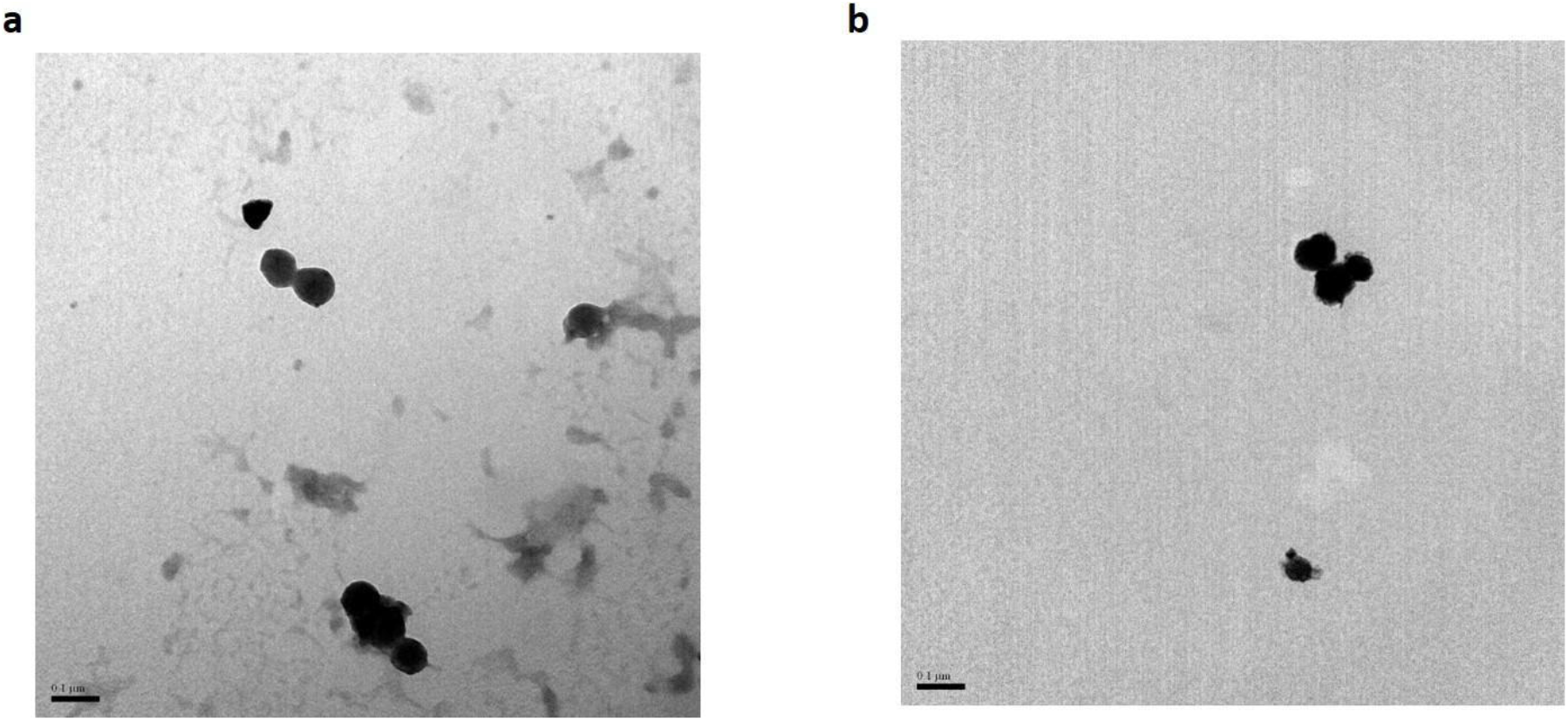
TEM analysis of Mn_3_O_4_ (a) and GNA35 (b) nanomaterials. Scale is shown on each of the panels.

Particle size and stability was determined as well through dynamic light scattering and z-potential, resuspended in ultrapure water and in human cells culture media (DMEM + 1% Bovine Fetal Serum (FBS)), at a concentration of 20 mg L^-1^. The observed average values for the NMs resuspended in water were significantly higher than those observed through TEM: 347±69 nm for Mn_3_O_4_ and 364±91 nm for GNA35, while their respective z-potential was −16,4 ± 0,9 and −20,9 ± 0,6. When the samples were resuspended in culture media, only GNA35 NMs were stable enough to perform DLS and z-potential measurements, which were very similar to those obtained in water (DLS: 338 ± 0,6 nm; z-potential: −23,0 ± 3,0).

The chemical composition and crystalline phase of Mn_3_O_4_ and GNA35 were determined by X-ray diffraction analysis. The assays were performed by X-Ray Polycrystalline Diffraction. Figure 2 shows the obtained diffraction patterns, which are characteristic of hausmannite (Mn_3_O_4_ reference pattern 00-024-0734), with a higher amount of impurities present in the raw material.

**Figure 2.**
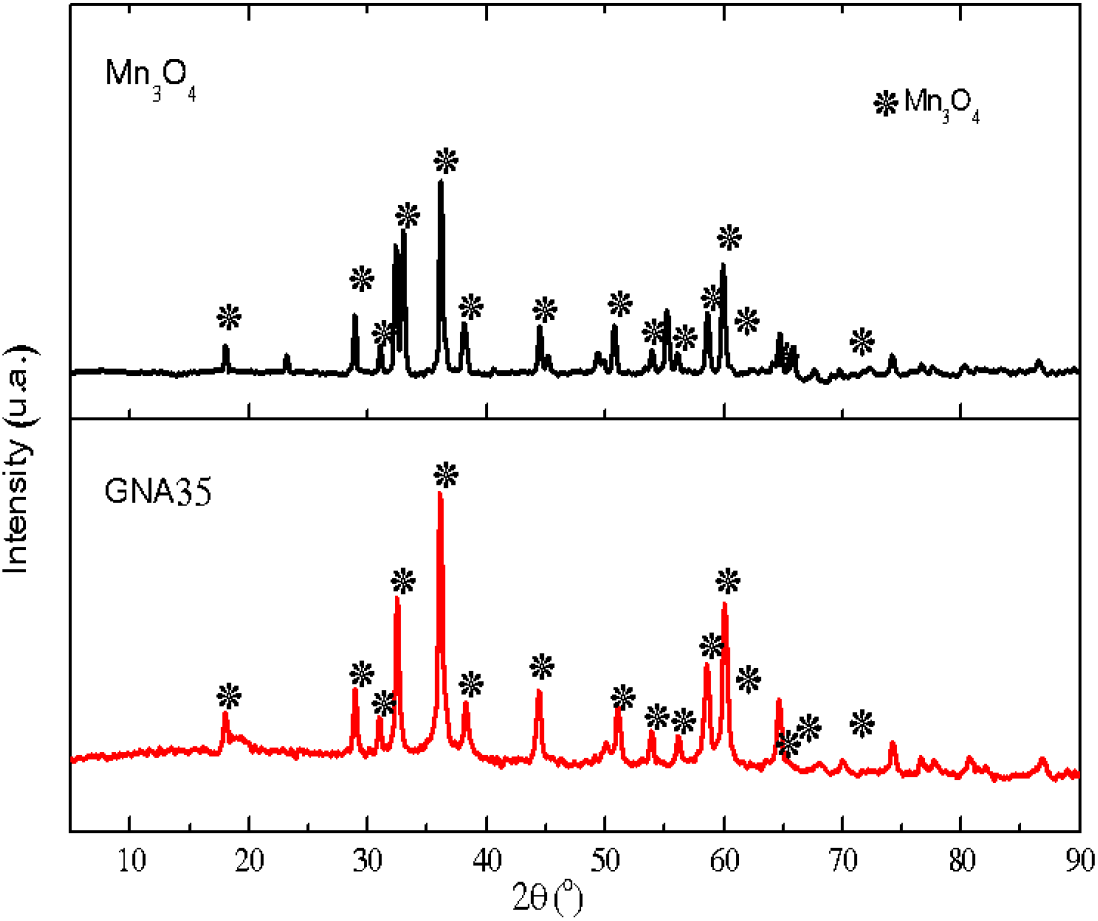
XRD spectra of Mn_3_O_4_ and GNA35 showing the diffraction peaks of hausmannite Mn_3_O_4_.

TGA analyses were carried out to determine the composition and the thermal stability of the manganese oxide nanoparticles. The obtained results are displayed in Figure 3, which indicate that the raw material presents an increase in weight probably due to an oxidation reaction, followed by a loss around 900 °C (Gillot et al., 2001; Amankwah and Pickles, 2009). However, GNA35 nanoparticle shows different weight losses with temperature variation due to the elimination of surface and occluded water and phase transformations of the oxide structure.

**Figure 3.**
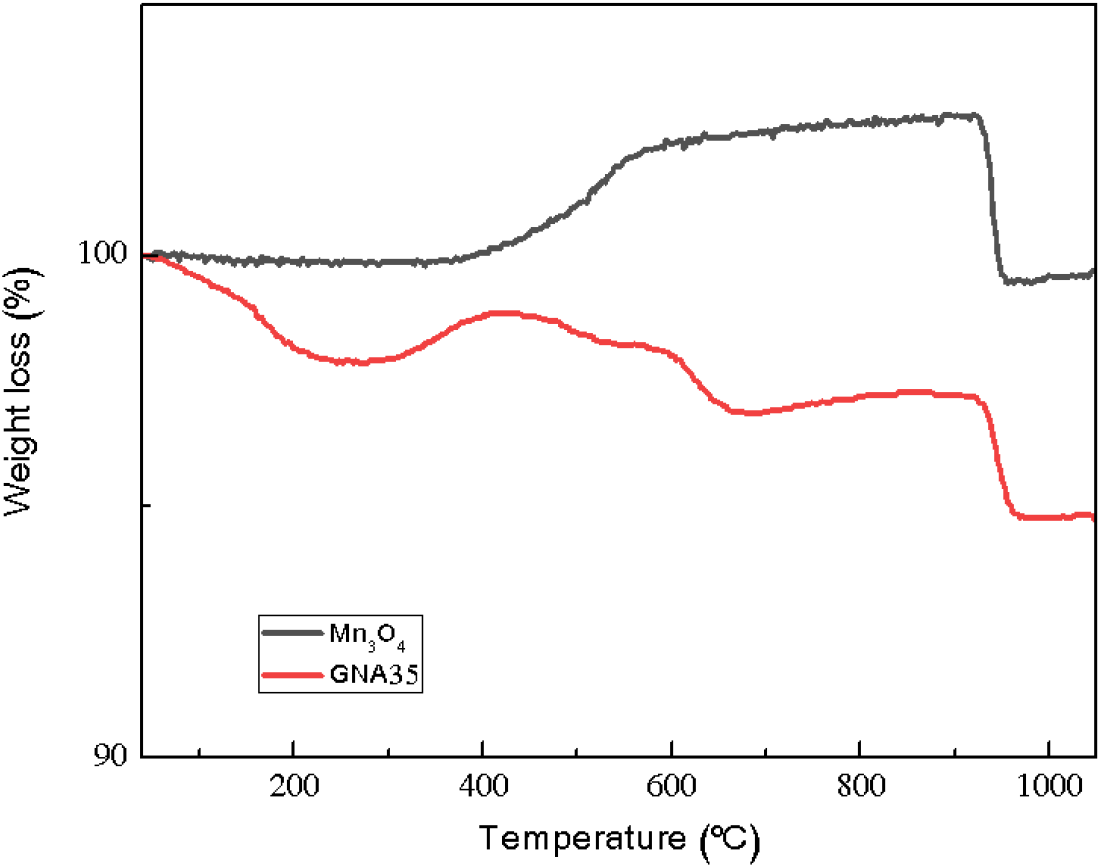
TG curves of Mn_3_O_4_ and GNA35. The composition of the nanoparticles was analyzed with the mass reduction as a function of temperature.

Analysis of nitrogen adsorption-desorption isotherms was carried out to determine the porosity of samples. The results have demonstrated a good agreement between surface area and accumulated charge, which is key in electrode materials for energy storage devices. The estimated BET area was 14 and 39 m^2^g^-1^ for Mn_3_O_4_ and GNA35 respectively, demonstrating that the synthesised material possesses increased specific surface, that could result in improved physical-chemical properties for charge accumulation.

Cyclic voltammetry was employed to determine the amount of charge that can be accumulated by Mn_3_O_4_ and GNA35. As it can be seen in Figure 4, the wider area observed for GNA35 indicates that the accumulated charge in this material is greater than in the raw material. Furthermore, the specific Capacitance (amount of stored charge per unit change in electric potential), calculated from galvanostatic charge/discharge measurements at 1 A g^-1^ for GNA35 was 150 F g^-1^, being 16 F g^-1^ for Mn_3_O_4_. These results corroborate that the synthesized nanoparticles have an improved electrochemical response when compared to the starting material, ensuring a better performance as electrode in an energy storage device.

**Figure 4.**
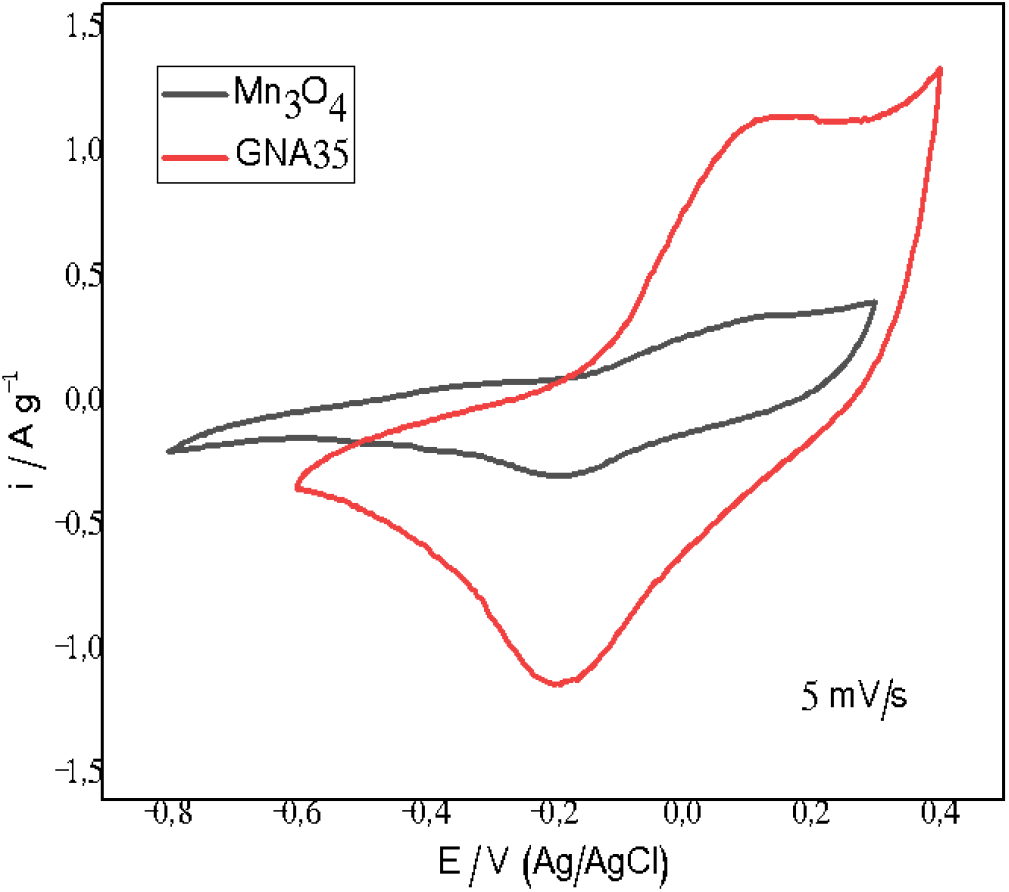
Cyclic voltametry of Mn_3_O_4_ and GNA35.

## 4. Toxicology assays

### 4.1 Determination of human alveolar carcinoma epithelial cell line (A549) response to GNA35 and Mn_3_O_4_

#### 4.1.1 MTT assay

The potential negative effect produced by the exposure of lung cells to the manganese oxide nanomaterials under study was assessed employing human alveolar carcinoma epithelial cells (A549) as model. The cytotoxicity of Mn_3_O_4_ and GNA35 on A549 cells was analysed using the MTT assay, exposing the cells to different concentrations of both nanomaterials (NMs) (1, 5, and 10 mg L^-1^) for 24 h. The obtained results indicate that both NMs are able to induce cytotoxic effects on the A549 cells at the selected concentrations, in a similar way, and in a concentration dependent manner (Figure 5). Cells incubated with Mn_3_O_4_ at 5 mg L^-1^ and 10 mg L^-1^ present a statistically significant reduction (*P* ≤ 0.05) in viability when compared to the non-exposed cells condition (control). In case of GNA35, the obtained results revealed statistically significant differences when cells were exposed to 1 mg L^-1^ (*P* ≤ 0.05), 5 mg L^-1^ (*P* ≤ 0.0001), and 10 mg L^-1^ (*P* ≤ 0.0001).

**Figure 5.**
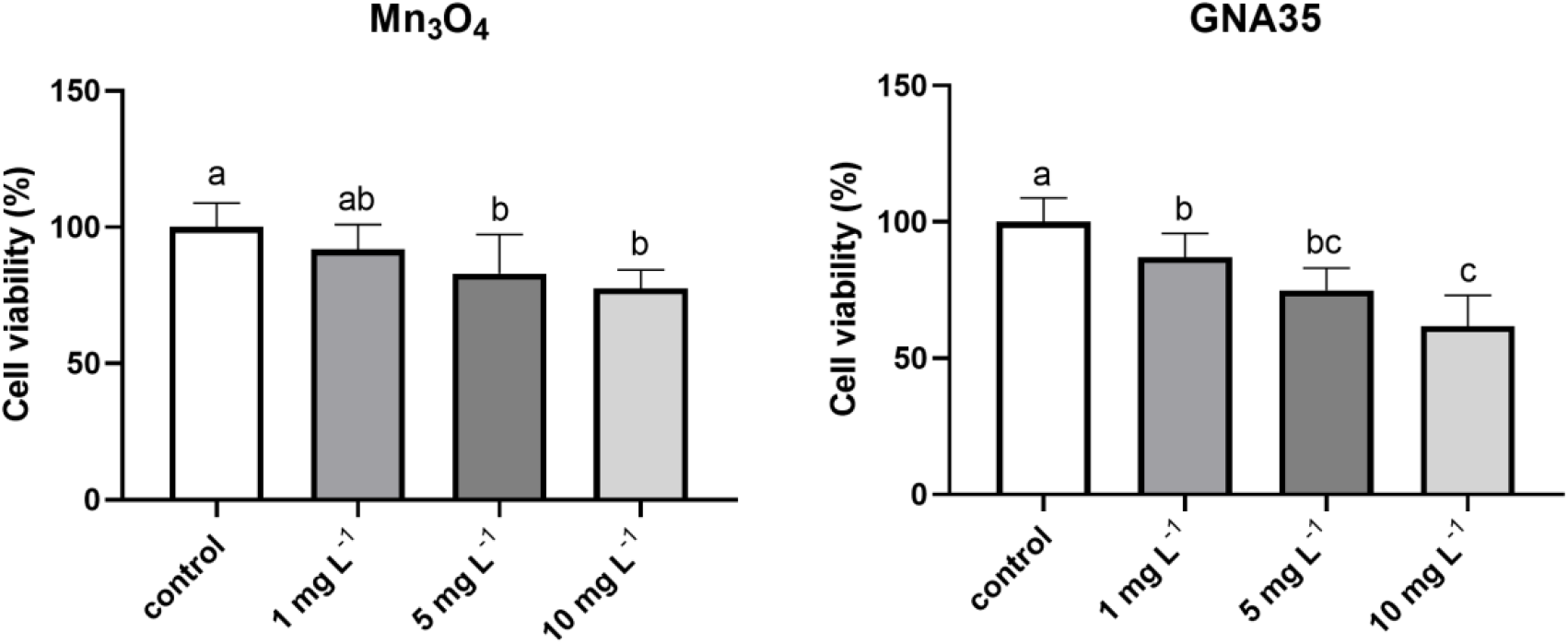
Viability of A549 cells exposed to different concentrations of Mn_3_O_4_ (**left**) and GNA35 (**right**). Results are expressed as % of control (untreated cells). Data represented the mean (± standard deviation, SD) of two independent experiments. Differences were established using a one-way ANOVA followed by multiple comparisons test (Tukey test) and considered significant when *P* ≤ 0.05. Different letters indicate significant differences between treatments.

#### 4.1.2 Oxidative stress assay

The induction of oxidative stress after A549 cells exposure to different Mn_3_O_4_ and GNA35 concentrations (1 and 10 mg L^-1^) was evaluated as well. The DCFH-DA assay was employed to determine the levels of reactive oxygen species (ROS) at different time points (0, 30 and 60 minutes), by measuring the increase of relative fluorescence. As displayed in Figure 6, ROS formation induced by both NMs could be observed, in function of time and of the concentration employed. In case of Mn_3_O_4_, the two selected concentrations induced an increased average of ROS levels in comparison with the non-exposed cells condition, but those were only significant in case of the higher concentration tested (*P* < 0,0001), 30 minutes after exposure. GNA35 induced ROS levels in A549 cells in a similar pattern to that observed for Mn_3_O_4_ NMs but triggering a stronger cellular response. In this case, the presence of 1 and 10 mg L^-1^ of GNA35 provoked an increase of relative fluorescence in A549 cells in a significant level (*P* < 0,001 and *P* < 0,0001 respectively), 30 minutes after exposure.

**Figure 6.**
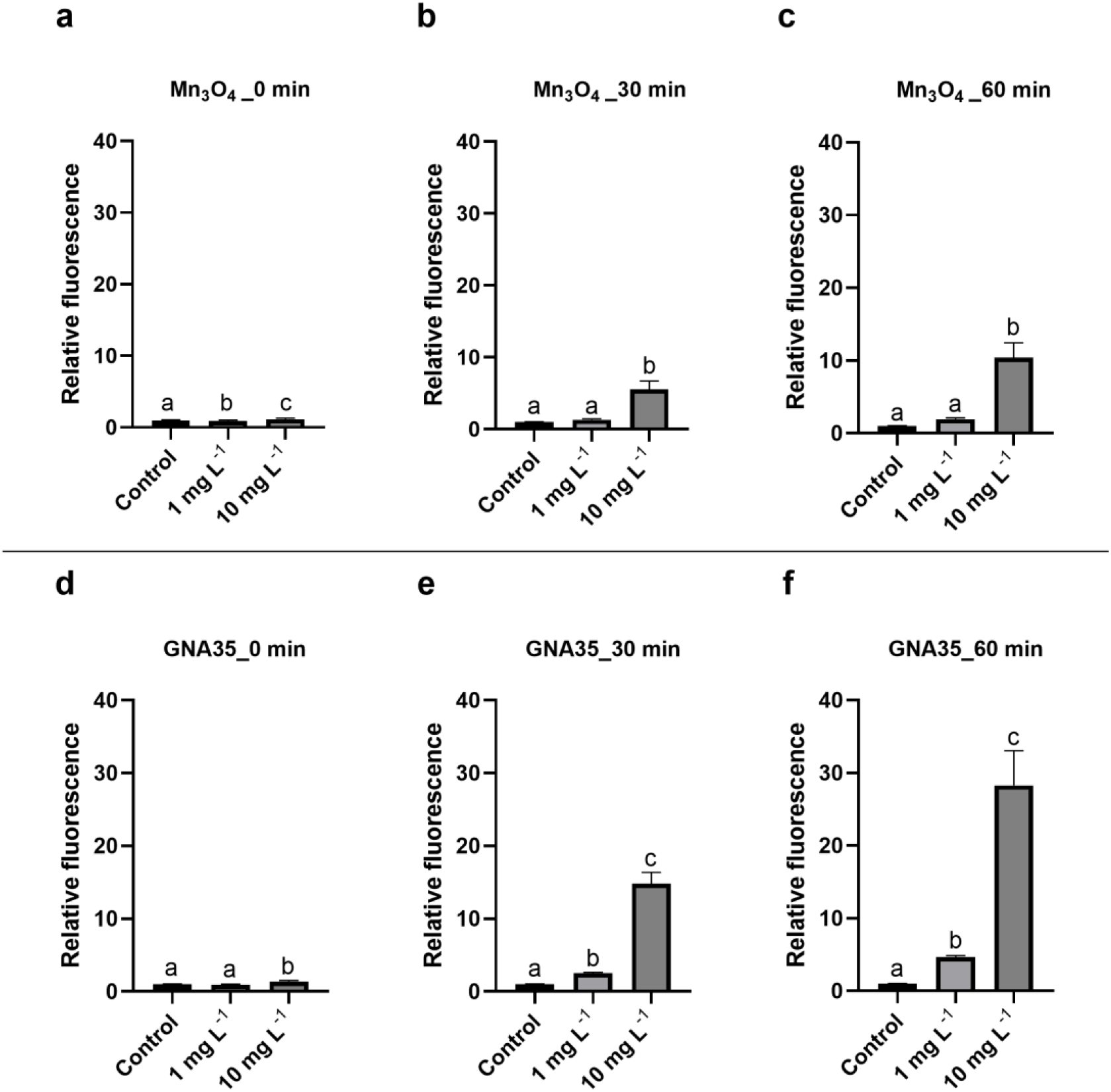
Comparison of the A549 ROS response against Mn_3_O_4_ (a-c) and GNA35 (d-f) nanoparticles at 0 minutes (a, d), 30 minutes (b, e) and 60 minutes (c, f). The charts have been represented at the same scale on the Y-axis. Differences were established using a one-way ANOVA followed by multiple comparisons test (Tukey test) and considered significant when *P* ≤ 0.05. Same letters show no significant differences between treatments.

### 4.2 Determination of human colon cancer cell line (HT29) response to GNA35 and Mn_3_O_4_

#### 4.2.1 MTT assay

The colon cancer cell line HT29 was selected as a model to predict the potential toxicity induced by manganese oxide nanomaterials upon exposure of human intestinal cells. The viability of the HT29 cell line was determined by MTT assay, after 24 h of exposure to Mn_3_O_4_ and GNA35 at 1, 5 and 10 mg L^-1^. In contrast to the observations made when A549 cells were exposed to the same concentrations of the selected NMs, HT29 cells did not present significant viability differences with the control condition, at any concentration tested (Figure 7).

**Figure 7.**
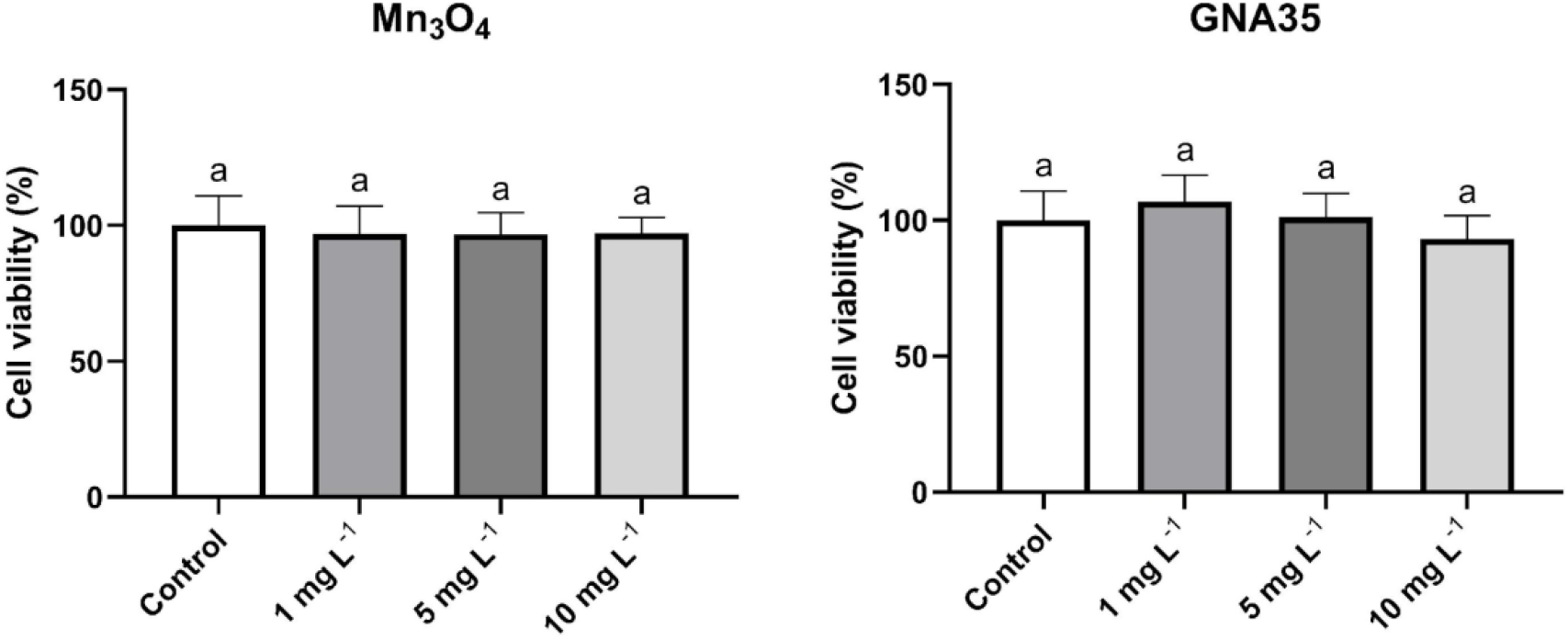
Viability of HT29 cells after the treatment with Mn_3_O_4_ (**left**) and GNA35 (**right**). Results are expressed as % of control (untreated cells). Data represented the mean (± standard deviation, SD) of two independent experiments. Differences were established using a one-way ANOVA followed by multiple comparisons test (Tukey test) and considered significant when *P* ≤ 0.05.

#### 4.2.2 Oxidative stress assay

The induction of oxidative stress after the exposure of HT29 cells by Mn_3_O_4_ and GNA35 was also assessed using the DCFH-DA assay, following the same exposure conditions as those employed with the A549 cell line. In regard to the obtained results, as it can be observed in Figure 8, both NMs provoked an increase in HT29 cells ROS levels, in a similar fashion to those induced in A549 cells. The increase in ROS levels could be directly associated to the employed concentrations and the exposure time. In case of Mn_3_O_4_, significantly higher ROS levels were measured in the presence of 1 and 10 mg L^-1^ after 30 minutes exposure (*P* < 0,001 and *P* < 0,0001 respectively). When evaluating GNA35, we found significant differences on relative fluorescence levels already at t0, when cells were exposed to 10 mg L^-1^ (*P* < 0,05), while no significant differences were observed in the presence of 1 mg L^-1^ at any of the measured timepoints. As described above for A549 cells, GNA35 also induced stronger ROS levels on HT29 cells in comparison to Mn_3_O_4_ NMs.

**Figure 8.**
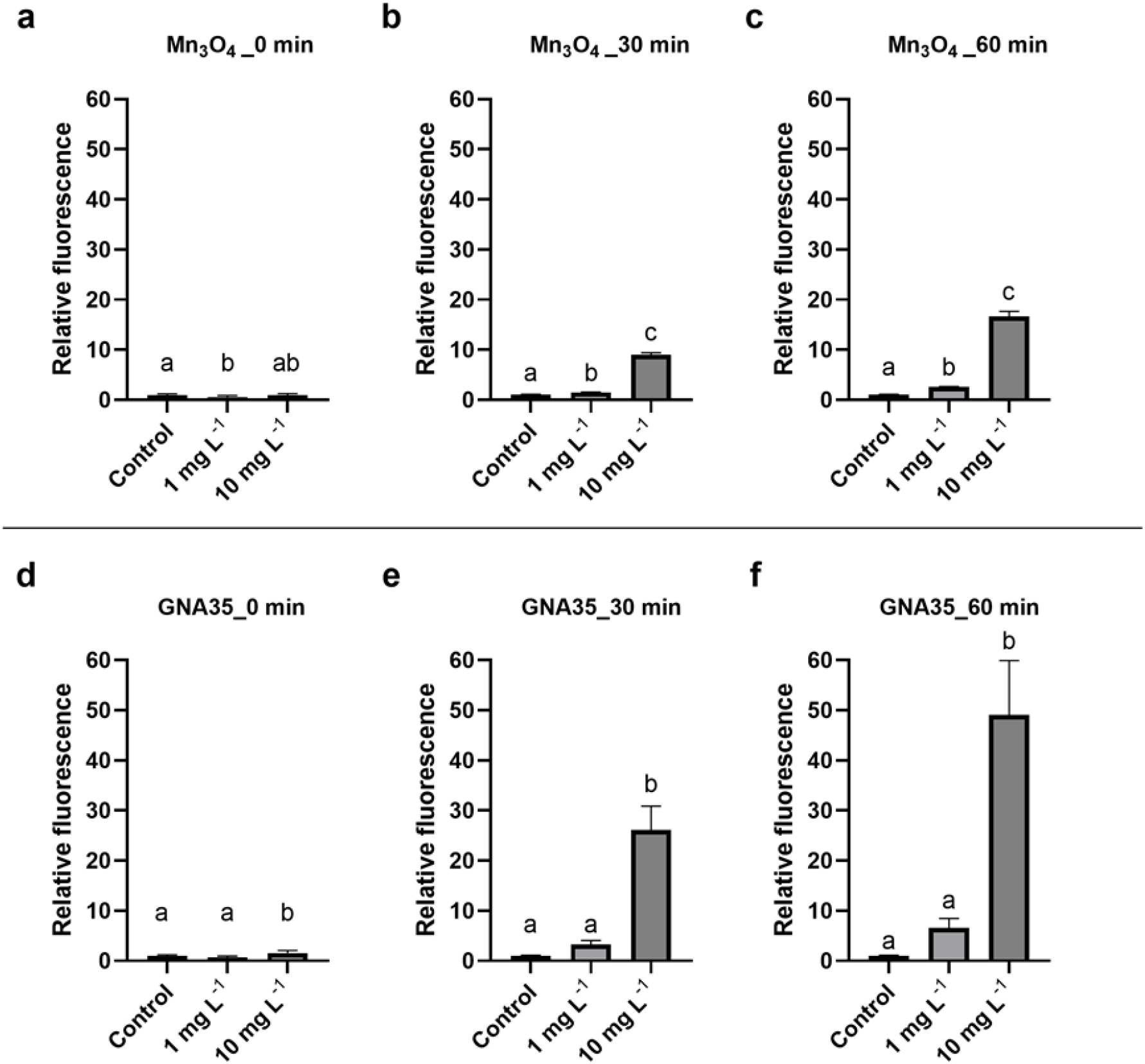
Comparison of the HT29 ROS response against Mn_3_O_4_ (a-c) and GNA35 (d-f) nanoparticles at 0 minutes (a, d), 30 minutes (b, e) and 60 minutes (c, f). The charts have been represented at the same scale on the Y axis. Same letters show no significant differences between treatments.

### 4.3 Determination of human reconstructed human epidermis (RhE) response to GNA35 and Mn_3_O_4_

#### 4.3.1 *In Vitro* EpiDerm^™^ Skin Irritation Test

The potential of Mn_3_O_4_ and GN35 to induce skin irritation was evaluated using the *in Vitro* EpiDerm^™^ Skin Irritation Test (EPI-200-SIT), following OECD guidelines (Test No. 439). As described in the corresponding Materials and Methods section, the procedure involves the use of reconstructed human epidermis (RhE), which consists of a 3D cell culture closely mimicking the biochemical and physiological properties of the upper parts of the human skin, employing the MTT assay to determine cell viability. According to EU and Globally Harmonized System of Classification and Labelling Chemicals, GHS, (R38/ Category 2 or no label), an irritant is predicted if the mean relative tissue viability of three individual tissues exposed to the test substance (in this case, the nanoparticles) is reduced below 50% of the mean viability of the negative controls. As it can be observed in Figure 9, a high concentration (500 mg L^-1^) of Mn_3_O_4_ and GNA35 did not reduce the viability of RhE. In contrast, the positive control (tissues exposed to 5% SDS) indicated by OECD TG439 reduced the RhE viability as expected (>95%). Therefore, both nanoparticles can be considered as non-irritant in the conditions tested.

**Figure 9.**
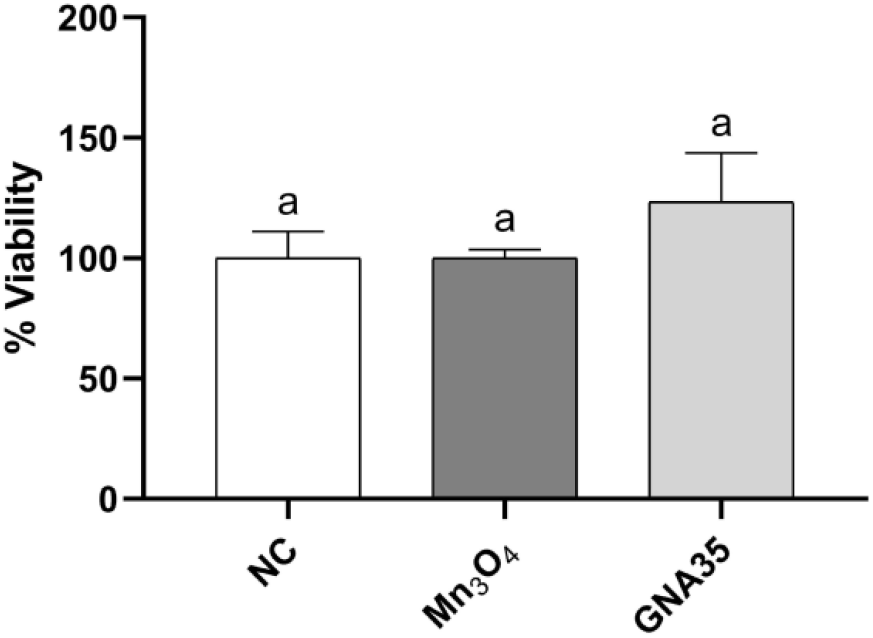
EpiDerm™ tissues were exposed to 500 mg L^-1^ of Mn_3_O_4_ and GN35 for 1 h. The viability was analysed by MTT assay, and it is expressed as a percent of negative control. Data represented the mean ± standard deviation (SD).

#### 4.3.2 Cytokine release assay

The concentration of IL-1α released from the RhE tissues into the assay medium during the exposure to Mn_3_O_4_ and GNA35 was measured by ELISA test and expressed as pg mL^-1^ released by RhE after each skin irritation test treatment. The IL-1α values obtained were 58.6 ± 34.0 pg mL^-1^ for the non-exposed cells, 146.6 ± 79.9 pg mL^-1^ for Mn_3_O_4_, and 69.7 ± 33.3 pg mL^-1^ for GNA35. Although the average IL-1α released in the presence of Mn_3_O_4_ was higher than in the rest of the conditions tested, the observed differences were not considered statistically significant.

## 5. Discussion

Nanomaterials (NMs) have brought a myriad of applications into material science due to an expanded set of interesting properties (Khan et al., 2016). Thus, their rapid commercialization and their widespread use has led to the need to study their effect on human health and the environment (Ganguly et al., 2018; Singh et al., 2019), especially from an occupational perspective. Toxicity of most of the available nanomaterials is unknown to a great extent, besides lacking of regulated occupational exposure limits (Pietroiusti et al., 2018). Their small size and large surface area facilitate their entry in the human body through different routes, which makes toxicological studies necessary to elucidate the potential adverse effects of the nanoparticles (Singh et al., 2013). Nanotoxicology is a novel interdisciplinary approach for the evaluation of the safety of these materials. This field is migrating from its infancy to a high degree of maturity, and the evaluation of toxic effects of nanoparticles are not limited just to the enumeration of damage and cell death, but it is approaching a state where it could provide useful insights to the manufacturers for the application of safe-by-design strategies for their materials. Besides, their workers can benefit from a first-hand occupational exposure hazard risks assessment. It is true that legislation has not taken enough steps in this direction, thus, the evaluation of material toxicity is of paramount importance for adequate policies to be taken into consideration.

The production and the use of Mn oxide NMs has increased in the last decades, leading to a major exposure and, consequently to a major risk for humans and the environment (Singh et al., 2013). In this work, two Mn oxide nanomaterials, Mn_3_O_4_ and GNA35, were studied. GNA35 NMs were designed to improve the electrochemical functionalities of the commercial Mn_3_O_4_, a necessary step for its use in applications such as the development of supercapacitators. The analysis of the physico-chemical properties of both NMs revealed similarities in terms of morphology and size, but differences in terms of stability, composition and electrochemical performance. TEM analysis showed that both materials are nanoparticles with round shape and a similar diameter, of around 100 nm or smaller. In both cases, DLS and z-potential analysis revealed a certain colloidal instability when the NMs were resuspended in ultrapure water, higher in case of Mn_3_O_4_. Stability differences between GNA35 and Mn_3_O_4_ were more noticeable when the NMs were resuspended in human cells culture media. In this case, GNA35 maintained a similar stability to that observed in ultrapure water, while Mn_3_O_4_ instability even prevented to perform accurate DLS and z-potential measurements. XRD and TGA analyses showed compositional differences between both NMs. Mn_3_O_4_ and GNA35 conserved the characteristic pattern of hausmannite, although the former showed to have a higher degree of impurities.

Surface area and cyclic voltammetry analyses were performed to understand the potential of the novel NM for its use in electrochemical applications. In both cases, GNA35 showed improved properties with respect to the precursor NM. In terms of surface area, which was measured by analyzing nitrogen adsorption-desorption isotherms, GNA35 (39 m^2^g^-1^) showed to have a higher estimated BET area than Mn_3_O_4_ (14 m^2^g^-1^). Cyclic voltammetry analysis allowed the calculation of the specific capacitance of the NMs, from galvanostatic charge/discharge measurements at 1 A g^-1^. The calculated specific capacitance for GNA35 (150 F g^-1^) was around one order of magnitude higher than that observed for Mn_3_O_4_ (16 F g^-1^). The obtained results demonstrated that GNA35 have significantly improved properties for charge accumulation when compared with the starting material, ensuring a better performance as electrode in an energy storage device.

Once the relevant physico-chemical properties of both NMs were understood, the toxicity of GNA35 and Mn_3_O_4_ was evaluated in different cellular models mimicking human tissues that could be directly exposed to them. There are numerous routes through which a chemical can enter the human body when considering, occupational or environmental exposures, including, inhalation, ingestion, injection or contact with the skin (Egorova and Ananikov, 2017). The way of entry is a very important factor, and for this reason, we have evaluated the potential toxic effects of the NMs over three different models suitable to perform a hazard assessment associated to potential inhalation, ingestion and dermal contact: two cell lines representative of lung and gastrointestinal system, and an *in vitro* skin model.

As mentioned previously, the respiratory system is the main pathway for airborne contaminant entry into the body, allowing them to be transported to the other organs (Sonwani et al., 2021). Considering this, we have analysed the cytotoxicity of Mn_3_O_4_ and GNA35 nanoparticles in the human alveolar carcinoma epithelial cell line (A549). The obtained results (viability determination and oxidative stress formation) showed a dose-dependent cytotoxicity within the employed concentration range (1-10 mg L^-1^). These results are in accordance with the work of Shaik *et al.,* who showed as well a decrease in cell viability on the range of concentrations over 10-25 mg L^-1^ in response to Mn_3_O_4_ nanoparticles exposure (Shaik et al., 2021). Interestingly, manganese oxide NMs coated with polyethylene glycol (PEG) have shown very low toxicity, in terms of cell viability reduction, towards the same cell line, in a higher concentration range (200-1000 mg L^-1^) (Zhan et al., 2017). In regard to oxidative stress induction, it is well known that metal oxide particles, including manganese oxide, induce ROS production, leading to cell damage (Alarifi et al., 2017). Specifically, the generation of particularly high ROS levels upon exposure to Mn_3_O_4_ has been reported (Limbach et al., 2007), which is in concordance with the results obtained in the present study.

Other relevant via of potential exposure to nanoparticles is through accidental ingestion. Their accumulation in the intestine can produce toxicity and, consequently, increase the risk of developing colon cancer or other carcinomas (Sonwani et al., 2021). In this context, we have studied the effect of Mn_3_O_4_ and GNA35 on the human colon cancer cell line (HT29). In contrast to the results obtained for A549 cells, the viability of HT29 cells was not reduced after the exposure to both nanoparticles at 1, 5, 10 mg L^-1^. Khan *et al.* made a different observation in a previous study, where the exposure of HT29 cells to Mn_3_O_4_ reduced their viability at 5 and 10 mg L^-1^ (Khan et al., 2016). However, relevant characteristics that could affect the potential toxicity of the employed NMs, such as size, colloidal stability or BET were not revealed, which makes difficult to compare the obtained results in both research works. In any case, Mn_3_O_4_. and GNA35 enhanced ROS levels of HT29 cells in a similar way to what was observed for A549 cells, which shows their cytotoxicity towards the colon cell line. In this regard, Choi *et al*. have demonstrated that manganese oxide NMs are able to induce the production of ROS in different human cell lines (Choi et al., 2010), indicating the ability of manganese oxide nanomaterials to generate cellular damage in human cells, regardless the type.

Skin contact is another relevant potential nanomaterial exposure route. It is the largest organ in the body (1.5-2 m^2^ of surface area), and it plays a very important role as a protective barrier against environmental allergens, pathogens, chemicals, and harmful materials (Wang et al., 2018). In particular, metallic nanoparticles can release ions that could cause sensitization and irritation after the exposure, but not enough data has been made available in this regard. For this reason, more transdermal toxicity studies are required to understand the adverse effects of metallic nanoparticles on the skin (Sufian et al., 2017; Hashempour et al., 2019). In the present work, we have studied the irritant potential of Mn_3_O_4_ and GNA35 nanoparticles on the reconstructed human epidermal model EpiDerm™. Following the OECD test n°439 to determine *in vitro* skin irritation on reconstructed human epidermis (RhE), we have observed that none of the NMs produce damage in the tissue. According to EU and Globally Harmonized System of Classification and Labelling Chemicals, GHS, (R38/ Category 2 or no label), an irritant is predicted if the mean relative tissue viability of three individual tissues exposed to the test substance is reduced below 50% of the mean viability of the negative controls. In case of Mn_3_O_4_ and GNA35 nanoparticles, they did not reduce the viability of the *in vitro* model. Considering this, both nanoparticles can be considered as non-irritant in the studied conditions. Regarding the possible induction of an inflammatory skin process by Mn_3_O_4_ and GNA35, we evaluated the IL-1α secretion of the exposed RhE. The obtained results showed that none of the NMs produce a significant increase in the release of the interleukin. So far, to our knowledge, there are no studies in the literature regarding the irritant or inflammatory potential of manganese oxide nanoparticles in RhE models. However, our results are in the same line with those described for other metallic nanoparticles (Fe NPs, Al NPs, Ti NPs, and Ag NPs), which resulted to be non-irritant and non-inflammatory when the same test method and *in vitro* skin model was employed (Park et al., 2011; Choi et al., 2014; Kim et al., 2016; Miyani and Hughes, 2017).

Overall, the obtained results indicate that both the precursor product (Mn_3_O_4_) and the synthesized nanomaterial with enhanced properties for its use in energy storage applications (GNA35) have similar hazard characteristics, inducing cytotoxicity effects in the selected lung and colon *in vitro* models, while lacking irritation potential in the skin model. In terms of the cytotoxicity effects observed, GNA35 induced the production of higher ROS levels than Mn_3_O_4_ in the A549 and HT29 human cell lines, which might be related to the higher specific surface and surface charge observed in GNA35. Both surface-related modes of action have been defined as key factors for nanomaterials toxicity (Fröhlich, 2012; Shin et al., 2015; Schmid and Stoeger, 2016).

## 6. Conclusion

The results presented in this research article reveal the physico-chemical properties and the potential hazard of novel manganese oxide NMs developed for their use in the manufacture of energy storage devices. While GNA35 physicochemical characteristics showed an enhanced surface area and electrochemical response in comparison with the precursor material (Mn_3_O_4_), the study of their toxicological properties towards human *in vitro* models representing three potential exposure routes (alveolar, intestinal and dermal) revealed a similar response in the selected exposure conditions. The manganese oxide nanomaterials under study were able to reduce A549 cells viability, while no effect in the HT29 cells viability was observed. Both NMs provoked oxidative stress in the lung and colon *in vitro* models. In both cases, GNA35 induced the production of higher ROS levels than Mn_3_O_4_ which might be related to the higher specific surface and surface charge observed in GNA35. In case of the dermal exposure, no effects were observed in the irritation and inflammation tests performed. Therefore, according to EU and Globally Harmonized System of Classification and Labelling Chemicals, Mn_3_O_4_ and GNA35 are not considered as irritants in the exposure conditions tested.

## 7. Acknowledgements

This project was funded by grant No. NANOCOMP - BU058P20, from Junta de Castilla y León (Spain). The funding source had no role in study design, sample or data collection, data analysis or interpretation, manuscript writing, in the conclusions, or in the decision to submit this study for publication.

## References

Alarifi, S., Ali, D., Alkahtani, S., and Almeer, R. S. (2017). ROS-Mediated Apoptosis and Genotoxicity Induced by Palladium Nanoparticles in Human Skin Malignant Melanoma Cells. Oxid. Med. Cell. Longev. 2017. doi:10.1155/2017/8439098.

Alhadlaq, H. A., Akhtar, M. J., and Ahamed, M. (2019). Different cytotoxic and apoptotic responses of MCF-7 and HT1080 cells to MnO2 nanoparticles are based on similar mode of action. Toxicology 411, 71–80. doi:10.1016/j.tox.2018.10.023.

Amankwah, R. K., and Pickles, C. A. (2009). Thermodynamic, thermogravimetric and permittivity studies of hausmannite (Mn3O4) in air. J. Therm. Anal. Calorim. 98, 849–853. doi:10.1007/s10973-009-0273-3.

Choi, J., Kim, H., Choi, J., Oh, S. M., Park, J., and Park, K. (2014). Skin corrosion and irritation test of sunscreen nanoparticles using reconstructed 3D human skin model. Environ. Health Toxicol. 29. doi:10.5620/eht.2014.29.e2014004.

Choi, J. Y., Lee, S. H., Na, H. Bin, An, K., Hyeon, T., and Seo, T. S. (2010). In vitro cytotoxicity screening of water-dispersible metal oxide nanoparticles in human cell lines. Biorocess Biosyst. Eng. 33, 21–30. doi:10.1007/s00449-009-0354-5.

Circu, M. L., and Aw, T. Y. (2010). Reactive oxygen species, cellular redox systems, and apoptosis. Free Radic. Biol. Med. 48, 749–762. doi:10.1016/j.freeradbiomed.2009.12.022.

Clarke, C., and Upson, S. (2017). A global portrait of the manganese industry—A socioeconomic perspective. Neurotoxicology 58, 173–179. doi:10.1016/j.neuro.2016.03.013.

Dawadi, S., Gupta, A., Khatri, M., Budhathoki, B., Lamichhane, G., and Parajuli, N. (2020). Manganese dioxide nanoparticles: synthesis, application and challenges. Bull. Mater. Sci. 43. doi:10.1007/s12034-020-02247-8.

Domi, B., Rumbo, C., García-Tojal, J., Sima, L. E., Negroiu, G., and Tamayo-Ramos, J. A. (2020). Interaction analysis of commercial graphene oxide nanoparticles with unicellular systems and biomolecules. Int. J. Mol. Sci. 21, 205. doi:10.3390/ijms21010205.

Egorova, K. S., and Ananikov, V. P. (2017). Toxicity of Metal Compounds: Knowledge and Myths. Organometallics 36, 4071–4090. doi:10.1021/acs.organomet.7b00605.

Frick, R., Müller-Edenborn, B., Schlicker, A., Rothen-Rutishauser, B., Raemy, D. O., Günther, D., et al. (2011). Comparison of manganese oxide nanoparticles and manganese sulfate with regard to oxidative stress, uptake and apoptosis in alveolar epithelial cells. Toxicol. Lett. 205, 163–172. doi:10.1016/j.toxlet.2011.05.1037.

Fröhlich, E. (2012). The role of surface charge in cellular uptake and cytotoxicity of medical nanoparticles. Int. J. Nanomedicine 7, 5577–5591. doi:10.2147/IJN.S36111.

Ganguly, P., Breen, A., and Pillai, S. C. (2018). Toxicity of Nanomaterials: Exposure, Pathways, Assessment, and Recent Advances. ACS Biomater. Sci. Eng. 4, 2237–2275. doi:10.1021/acsbiomaterials.8b00068.

Ghosh, T., Zhang, W., Ghosh, D., and Kechris, K. (2020). ‘Predictive Modeling for Metabolomics Data’, in Computational Methods and Data Analysis for Metabolomics Methods in Molecular Biology., ed. S. Li (New York, NY: Springer US), 313–336.

Gillot, B., Guendouzi, M. El, and Laaij, M. (2001). Particle size effects on the oxidation–reduction behavior of Mn3O4 hausmannite. 70, 54–60.

Hashempour, S., Ghanbarzadeh, S., Maibach, H. I., Ghorbani, M., and Hamishehkar, H. (2019). Skin toxicity of topically applied nanoparticles. Ther. Deliv. 10, 383–396. doi:10.4155/tde-2018-0060.

Khan, S., Ansari, A. A., Khan, A. A., Abdulla, M., Al-Obeed, O., and Ahmad, R. (2016). In vitro evaluation of anticancer and biological activities of synthesized manganese oxide nanoparticles. Medchemcomm 7, 1647–1653. doi:10.1039/c6md00219f.

Kim, H., Choi, J., Lee, H., Park, J., Yoon, B.-I., Jin, S. M., et al. (2016). Skin Corrosion and Irritation Test of Nanoparticles Using Reconstructed Three-Dimensional Human Skin Model, EpiDerm^TM^. Toxicol. Res. 32, 311–316. doi:10.5487/TR.2016.32.4.311.

Limbach, L. K., Wick, P., Manser, P., Grass, R. N., Bruinink, A., and Stark, W. J. (2007). Exposure of engineered nanoparticles to human lung epithelial cells: Influence of chemical composition and catalytic activity on oxidative stress. Environ. Sci. Technol. 41, 4158–4163. doi:10.1021/es062629t.

Lucchini, R. G., Dorman, D. C., Elder, A., and Veronesi, B. (2012). Neurological impacts from inhalation of pollutants and the nose-brain connection. Neurotoxicology 33, 838–841. doi:10.1016/j.neuro.2011.12.001.

Martinez-Finley, E. J., Gavin, C. E., Aschner, M., and Gunter, T. E. (2013). Manganese neurotoxicity and the role of reactive oxygen species. Free Radic. Biol. Med. 62, 65–75. doi:10.1016/j.freeradbiomed.2013.01.032.

Miyani, V. A., and Hughes, M. F. (2017). Assessment of the in vitro dermal irritation potential of cerium, silver, and titanium nanoparticles in a human skin equivalent model. Cutan. Ocul. Toxicol. 36, 145–151. doi:10.1080/15569527.2016.1211671.

Nádaská, G., Lesný, J., and Michalík, I. (2010). Environmental aspect of manganese chemistry. Hungarian J. Sci. ENV-100702-A, 1–16.

Park, Y.-H., Jeong, S. H., Yi, S. M., Choi, B. H., Kim, Y.-R., Kim, I.-K., et al. (2011). Analysis for the potential of polystyrene and TiO2 nanoparticles to induce skin irritation, phototoxicity, and sensitization. Toxicol. Vitr. 25, 1863–1869. doi:10.1016/j.tiv.2011.05.022.

Peng, T., Xu, L., and Chen, H. (2010). Preparation and characterization of high specific surface area Mn3O4 from electrolytic manganese residue. Open Chem. 8, 1059–1068. doi:10.2478/s11532-010-0081-4.

Pietroiusti, A., Stockmann-Juvala, H., Lucaroni, F., and Savolainen, K. (2018). Nanomaterial exposure, toxicity, and impact on human health. WIREs Nanomedicine and Nanobiotechnology 10, e1513. doi:10.1002/wnan.1513.

Post, J. E. (1999). Manganese oxide minerals: Crystal structures and economic and environmental significance. Proc. Natl. Acad. Sci. U. S. A. 96, 3447–3454. doi:10.1073/pnas.96.7.3447.

Roels, H., Meiers, G., Delos, M., Ortega, I., Lauwerys, R., Buchet, J. P., et al. (1997). Influence of the route of administration and the chemical form (MnCl2, MnO2) on the absorption and cerebral distribution of manganese in rats. Arch. Toxicol. 71, 223–230. doi:10.1007/s002040050380.

Schmid, O., and Stoeger, T. (2016). Surface area is the biologically most effective dose metric for acute nanoparticle toxicity in the lung. J. Aerosol Sci. 99, 133–143. doi:10.1016/j.jaerosci.2015.12.006.

Shaik, M. R., Syed, R., Adil, S. F., Kuniyil, M., Khan, M., Alqahtani, M. S., et al. (2021). Mn3O4 nanoparticles: Synthesis, characterization and their antimicrobial and anticancer activity against A549 and MCF-7 cell lines. Saudi J. Biol. Sci. 28, 1196–1202. doi:10.1016/j.sjbs.2020.11.087.

Shin, S. W., Song, I. H., and Um, S. H. (2015). Role of physicochemical properties in nanoparticle toxicity. Nanomaterials 5, 1351–1365. doi:10.3390/nano5031351.

Singh, A. V., Laux, P., Luch, A., Sudrik, C., Wiehr, S., Wild, A.-M., et al. (2019). Review of emerging concepts in nanotoxicology: opportunities and challenges for safer nanomaterial design. Toxicol. Mech. Methods 29, 378–387. doi:10.1080/15376516.2019.1566425.

Singh, S. P., Kumari, M., Kumari, S. I., Rahman, M. F., Mahboob, M., and Grover, P. (2013). Toxicity assessment of manganese oxide micro and nanoparticles in Wistar rats after 28 days of repeated oral exposure. J. Appl. Toxicol. 33, 1165–1179. doi:10.1002/jat.2887.

Sonwani, S., Madaan, S., Arora, J., Suryanarayan, S., Rangra, D., Mongia, N., et al. (2021). Inhalation Exposure to Atmospheric Nanoparticles and Its Associated Impacts on Human Health: A Review. Front. Sustain. Cities 3, 1–20. doi:10.3389/frsc.2021.690444.

Sufian, M. M., Khattak, J. Z. K., Yousaf, S., and Rana, M. S. (2017). Safety issues associated with the use of nanoparticles in human body. Photodiagnosis Photodyn. Ther. 19, 67–72. doi:10.1016/j.pdpdt.2017.05.012.

Takeda, A. (2003). Manganese action in brain function. Brain Res. Rev. 41, 79–87. doi:10.1016/S0165-0173(02)00234-5.

Wang, M., Lai, X., Shao, L., and Li, L. (2018). Evaluation of immunoresponses and cytotoxicity from skin exposure to metallic nanoparticles. Int. J. Nanomedicine 13, 4445–4459. doi:10.2147/IJN.S170745.

Wang, P., Liang, C., Zhu, J., Yang, N., Jiao, A., Wang, W., et al. (2019). Manganese-Based Nanoplatform As Metal Ion-Enhanced ROS Generator for Combined Chemodynamic/Photodynamic Therapy. ACS Appl. Mater. Interfaces 11, 41140–41147. doi:10.1021/acsami.9b16617.

Zhan, Y., Zhan, W., Li, H., Xu, X., Cao, X., Zhu, S., et al. (2017). In vivo dual-modality fluorescence and magnetic resonance imaging-guided lymph node mapping with good biocompatibility manganese oxide nanoparticles. Molecules 22, 2208. doi:10.3390/molecules22122208.

Zhu, J., Wu, Q., and Li, J. (2020). Review And Prospect of Mn3O4-Based Composite Materials For Supercapacitor Electrodes. ChemistrySelect 5, 10407–10423. doi:10.1002/slct.202002544.

Zoni, S., Bonetti, G., and Lucchini, R. (2012). Olfactory functions at the intersection between environmental exposure to manganese and Parkinsonism. J. Trace Elem. Med. Biol. 26, 179–182. doi:10.1016/j.jtemb.2012.04.023.

